# Foxd3 controls heterochromatin-mediated silencing of repeat elements in mouse embryonic stem cells and represses the 2-cell transcription program

**DOI:** 10.1101/2021.05.01.442081

**Authors:** Deepika Puri, Birgit Koschorz, Bettina Engist, Megumi Onishi-Seebacher, Devon Ryan, Thomas Montavon

## Abstract

Repeat element transcription plays a vital role in early embryonic development. Expression of repeats such as MERVL characterises mouse embryos at the 2-cell stage, and defines a 2-cell-like cell (2CLC) population in a mouse embryonic stem cell culture. Repeat element sequences contain binding sites for numerous transcription factors. We identify the forkhead domain transcription factor FOXD3 as a regulator of repeat element transcription in mouse embryonic stem cells. FOXD3 binds to and recruits the histone methyltransferase SUV39H1 to MERVL and major satellite repeats, consequentially repressing the transcription of these repeats by the establishment of the H3K9me3 heterochromatin modification. Notably, depletion of FOXD3 leads to the de-repression of MERVL and major satellite repeats as well as a subset of genes expressed in the 2-cell state, shifting the balance between the stem cell and 2-cell like population in culture. Thus, FOXD3 acts as a negative regulator of repeat transcription, ascribing a novel function to this transcription factor.

## Introduction

Repetitive elements including tandem repeats and interspersed repeats constitute up to 45% of the mouse genome (Biémont, 2010). In most somatic cells, repeat elements are repressed by a combination of epigenetic modifications such as DNA methylation, and histone modifications such as H3K9me3, H4K20me3 and H3K27me3 (Peters *et al*, 2001; Martens *et al*, 2005; Kato *et al*, 2007; Mikkelsen *et al*, 2007). In early embryonic development however, these repeat elements are de-repressed. Activation of specific repeats along with thousands of zygotic genes, accompanied by the clearance of maternal transcripts constitutes an essential transcriptional milestone in embryonic development, called zygotic gene activation (ZGA) (Jukam *et al*, 2017). Pericentric major satellite repeats (MSRs) and the endogenous retrovirus MERVL are expressed at the 2-cell embryonic stage and contribute significantly to ZGA and normal embryonic development (Probst *et al*, 2010; Macfarlan *et al*, 2012; Burton & Torres-Padilla, 2014; Ishiuchi *et al*, 2015; Dang-Nguyen & Torres-Padilla, 2015). Additionally, mouse embryonic stem cells (mESCs), that have the property of self-renewal and differentiation into all three germ layers, also exist as a heterogeneous population in culture, consisting of a small fraction (1-5%) of cells that resemble the more totipotent 2C-like cells (2CLCs) (Macfarlan *et al*, 2012). These cells are characterised by an expanded potency and activation of MSRs and MERVL repeats along with a specific set of genes such as the Zscan4 family, Zfp352, Pramel7 etc. (Macfarlan *et al*, 2012; Genet & Torres-Padilla, 2020). Studies have shown that MERVL activation is sufficient for the conversion of mESCs to 2CLCs (Yang *et al*, 2020). During embryonic development, downregulation of MSR and MERVL is concomitant with increased LINE1 transcription, which facilitates the exit from the 2-cell state and contributes to ES cell self-renewal (Percharde *et al*, 2018). The importance of repeat element transcription in early development and stem cell function underscores the need to understand the transcriptional regulation of these repeats. Previous studies have demonstrated the presence of numerous transcription factor (TF) binding sites within repeat element sequences (Bourque *et al*, 2008; Bulut-Karslioglu *et al*, 2012). Transposable elements have emerged as a hub for transcription factor binding and assembly of transcription complexes (Hermant & Torres-Padilla, 2021). For example, PAX3 and PAX9 bind to and repress MSRs in mouse embryonic fibroblasts and this repression is essential for maintaining the integrity of heterochromatin (Bulut-Karslioglu *et al*, 2012). REX1 represses endogenous retroviruses (ERVs) in mESCs and preimplantation embryos through the binding and recruitment of YY1 and YY2 (Guallar *et al*, 2012). In contrast, ZSCAN4 and DUX act as positive regulators of MERVL transcription in mESCs and early embryos (Hendrickson *et al*, 2017; Zhang *et al*, 2019). Human ERVH and ERVK sequences contain binding sites for pluripotency TFs OCT4 and SOX2 which facilitate ERV transcription (Kunarso *et al*, 2010; Fort *et al*, 2014).Whether other transcription factors contribute to repeat regulation remains unknown.

This study focuses on the identification and characterisation of novel transcription factors that influence repeat element expression. Our analysis identifies the forkhead domain containing transcription factor FOXD3 as a novel regulator of repeat elements. FOXD3 is crucial for maintaining ES cell pluripotency (Hanna *et al*, 2002; Liu & Labosky, 2008). In stem cells, FOXD3 plays a bimodal role as an activator or a repressor in a context-dependent manner by enhancer decommissioning and recruitment of chromatin modulators such as the histone demethylase LSD1, chromatin remodelling factor BRG1 and histone deacetylases (HDACs) to target sites (Krishnakumar *et al*, 2016; Respuela *et al*, 2016; Sweet, 2016). We observe that in mESCs, FOXD3 binds to and represses MERVL and to a lesser extent, MSRs. Depletion of FOXD3 leads to a significant de-repression of MERVL and MSRs and a concomitant increase in a subset of 2CLC genes. Mechanistically, FOXD3 represses MERVL and MSRs by interacting with and recruiting the heterochromatin histone methyltransferase SUV39H1, which establishes the repressive H3K9me3 mark to the target sites. This study is a novel report of FOXD3 as a heterochromatin-mediated repressor of repeat element transcription and 2CLC gene expression in mouse embryonic stem cells.

## Materials and Methods

### Cell Culture

Foxd3 cKO cells obtained from Patricia Labosky lab were cultured using standard protocols on 0.2% gelatin coated plates. To induce Foxd3 knockout, cells were treated with 2μM 4-hydroxytamoxifen (Sigma-T5648) changed daily. Cells were harvested for experiments on day 2 when Foxd3 was depleted at RNA and protein level.

### Generation of recombinant GST-FOXD3 and GFP-FOXD3 fusion proteins

Wild type full-length Foxd3 cDNA was obtained from origene (Catalog number MR222218) and amplified using Foxd3 specific primers to be subcloned into the pGEX-6P1 plasmid and verified by sequencing. M1 (YSY-RAD) and M2 (FVK-VAM) mutants were generated by site directed mutagenesis using specific primers. GST-fusion proteins were expressed using protocols described in (Velazquez Camacho *et al*, 2017). Sequences encoding full length Foxd3, M1 and M2 mutants were subcloned into the pCAGGS-EGFP vector as described in (Velazquez Camacho *et al*, 2017). These plasmids were transfected into Foxd3 cKO cells using Xfect transfection reagent (Clontech). The cells were kept under puromycin selection to obtain polyclonal cell lines. All cell lines were checked periodically for mycoplasma contamination.

### EMSA

35 nucleotide 5’-Cy5 labelled and HPLC-purified DNA oligonucleotides were purchased from Sigma. To generate dsDNA, equimolar amounts of forward and reverse ssDNA oligonucleotides were mixed in 1xSSC buffer (150 mM NaCl, 15 mM sodium citrate) and incubated for 2 minutes at 90°C in a Thermomixer. The temperature was reduced to 60°C for 5 minutes, then further reduced to 20°C for 30 minutes. For EMSA, 50 nM of nucleic acids were mixed with increasing concentrations of recombinant proteins in a buffer containing 20 mM Tris-HCl pH 8.0, 100 mM KCl, 3mM MgCl2, 1 mM EDTA pH 8.0, 5% glycerol, 0.05% NP40, 2 mM DTT, 50 ng/ml yeast tRNA (ThermoFisher) and 2.5 ng/ml BSA (NEB). Samples were incubated at 4°C with rotation for 1 hour and resolved on a 4% polyacrylamide (60:1) gel (25 mM Tris-HCl, 200 mM glycine, 5% glycerol, 0.075% APS, 0.05% TEMED) in 12.5 mM Tris-HCl and 100 mM glycine. The Cy5 signal was scanned on a Typhoon FLA 9500 fluorescence scanner.

### Chromatin immunoprecipitation (ChIP)

ChIP was performed as described in (Bulut-Karslioglu *et al*, 2014) with antibody specific optimizations. For histone modification as well as FOXD3 ChIP, trypsinized cells were fixed by incubating with 1% formaldehyde for 10 minutes. For SUV39H1 ChIP, trypsinized cells were incubated in DSG ((Di (N-succinimidyl glutarate) (Synchem OHG)) at a final concentration of 2mM for 45 min at room temperature, washed twice with PBS and then incubated in 1% formaldehyde for 20 minutes at room temperature. Subsequent steps were performed as described in (Bulut-Karslioglu *et al*, 2014). The purified DNA was analysed by qPCR using specific primers as described in Table S2. The antibodies used for ChIP are as follows: Foxd3: Merck Millipore, Catalog number AB5687, 4μg. H3K9me3: crude serum, Antibody no 4861 (Jenuwein lab), 5μl. H3K4me3: Diagenode, Catalog number C15410003-50, 2μg. H3K27me3: Antibody no 6523 (Jenuwein lab) 4 μg, Suv39h1: Sigma Aldrich (Merck), Catalog number 05-615, 10μl.

### RNA Seq and analysis

Total RNA was extracted from Control and Foxd3 KO cells with Trizol (Invitrogen) and DNA was digested with Turbo DNase (Ambion), followed by clean-up with RNeasy MinElute Cleanup kit (Qiagen, 74204). Libraries were prepared using the TruSeq Stranded Total RNA Library Prep Gold (Illumina, 20020598) following Illumina protocols. The libraries were sequenced on a HiSeq2500 Illumina platform using a 100bp paired end approach. Two biological replicates were sequenced per cell line. The sequencing reads were aligned to the mouse genome build mm10 using TopHat2 (Kim *et al*, 2013) with default parameters. Repeats and genes were quantified using TEtranscripts (Jin *et al*, 2015) and DEseq2 (Love *et al*, 2014) was used to determine differentially expressed repeat elements and genes. deepTools was used to construct BigWig files and coverage tracks were visualized using IGV (Robinson *et al*, 2011). Detailed parameters are mentioned in (Velazquez Camacho *et al*, 2017). Heat map represented in figure 2f was plotted using Morpheus: https://software.broadinstitute.org/morpheus

**Figure 1:**
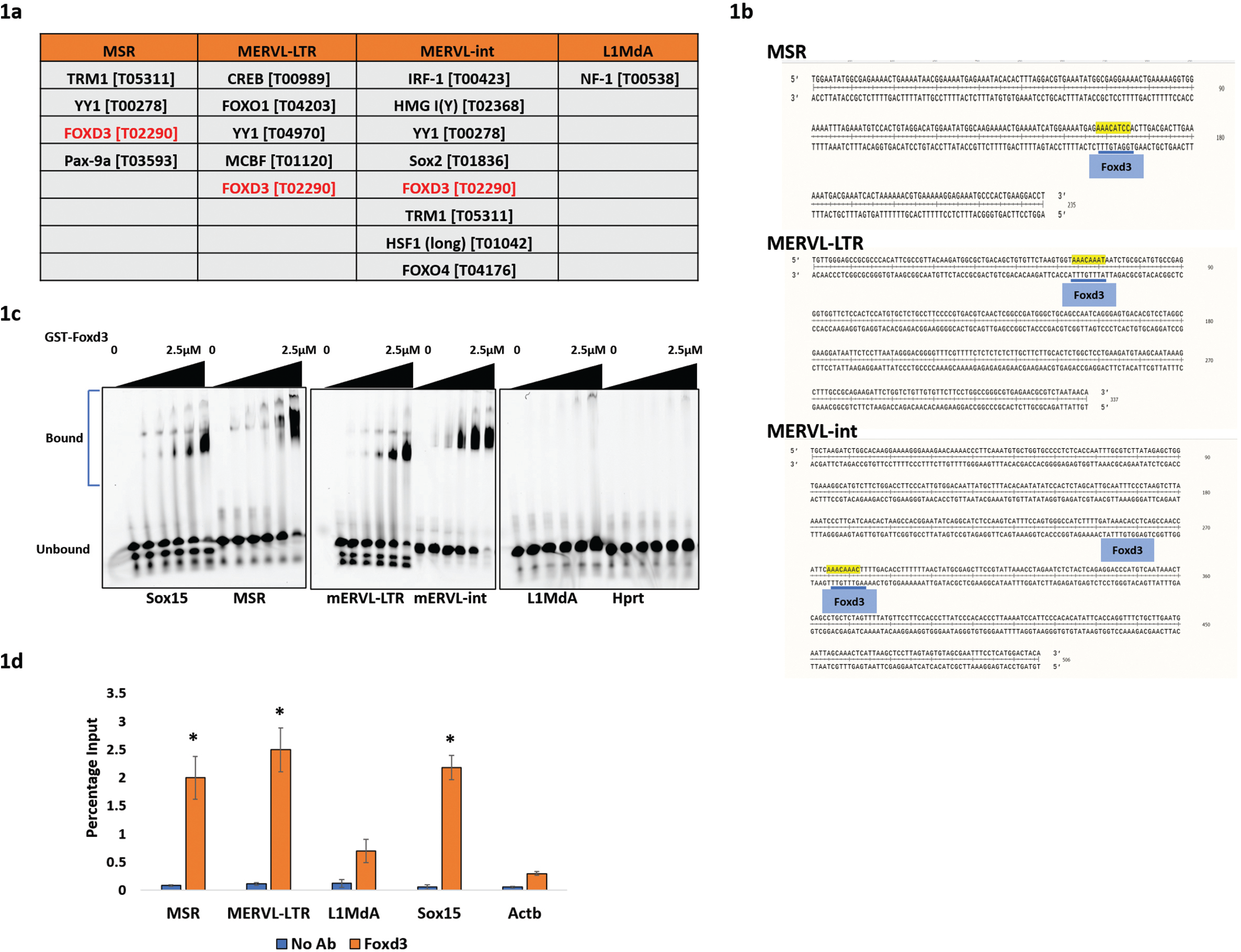
FOXD3 binds to MERVL and MSR elements. Figure 1a: Transcription factor binding sites with Transfac IDs predicted by PROMO in repeat element consensus sequences. Figure 1b: Consensus sequences of MSR, MERVL-LTR, MERVL-int with FOXD3 binding site highlighted (Blue boxes). The underlined sequences indicate statistically significant FOXD3 binding sites. Figure 1c: Electrophoretic mobility shift assay (EMSA) with increasing concentration (0, 15nM, 30nM, 60nM, 1.25μM and 2.5μM) of GST-FOXD3 and fixed concentration (50nM) of 5’-Cy5 labelled dsDNA oligonucleotides (35 bp each) from *Sox15* promoter (positive control), MSR, MERVL-LTR, MERVL-int, L1MdA and *Hprt* promoter (negative control). Figure 1d: ChIP-qPCR enrichment of FOXD3 in mESCs using primers specific for MERVL, MSR, L1MdA sequences. *Sox15* promoter and *Actb* promoter primers are used as positive and negative control respectively. Data is represented as percentage of input and the average of 3 biological replicates is plotted. Error bar indicates standard error of the mean (SEM). Asterisks indicate statistically significant differences compared to no antibody control levels (*: p< 0.05, paired t-test).

**Figure 2:**
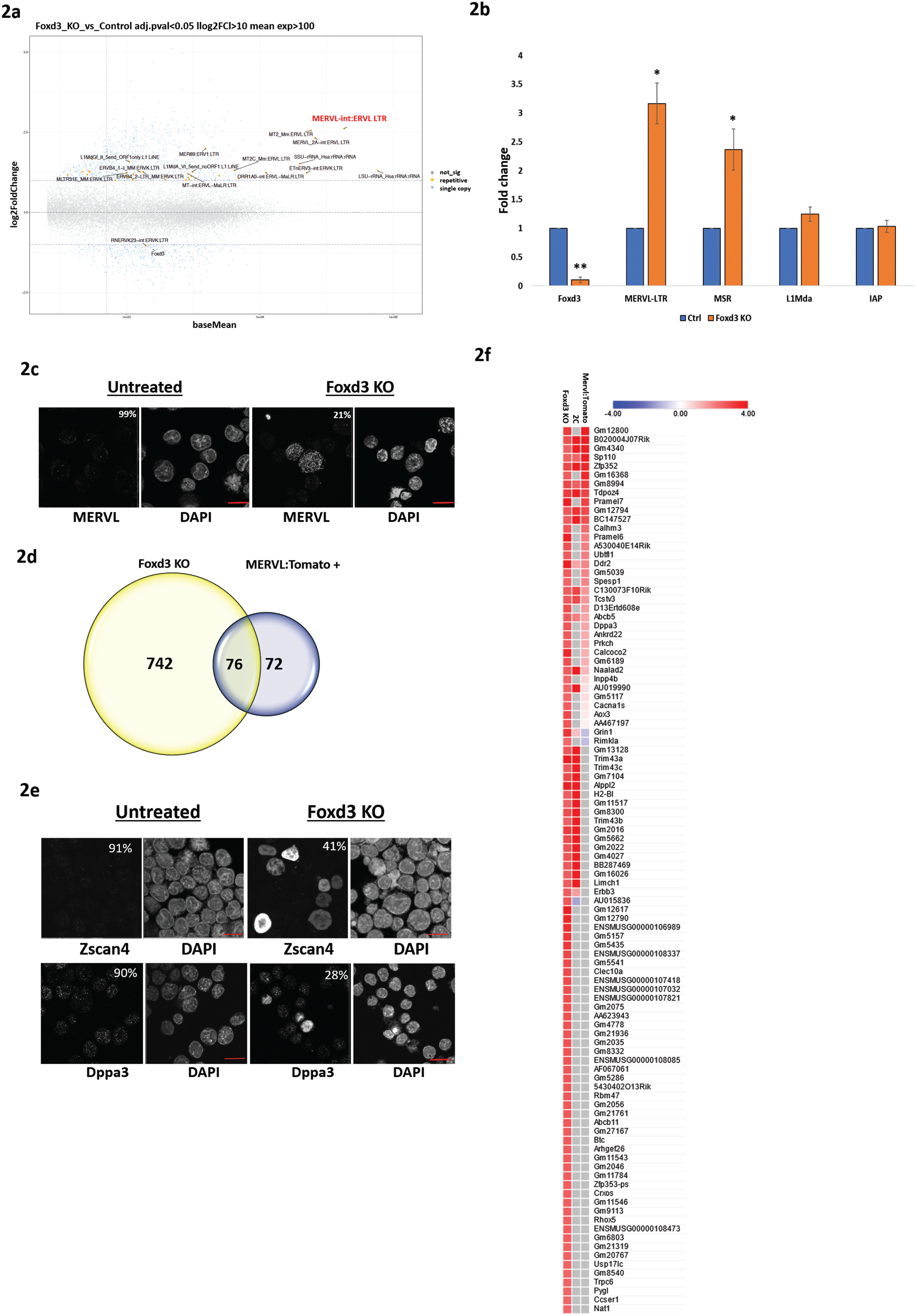
MSR, MERVL and 2CLC specific genes are upregulated in Foxd3 KO cells: Figure 2a: MA plot showing analysis of up and down regulated genes and repeats in total RNA-Seq preparations of control and Foxd3 KO cells. Significantly dysregulated genes are shown in blue and repeats are shown in yellow. The most highly upregulated repeat is marked in red. Figure 2b: Validation of select upregulated repeats in Foxd3 KO cells by RT-qPCR. The data is plotted as average fold change relative to control, after normalization to *Gapdh*. Error bars indicate SEM (N=3 biological replicates). Asterisks indicate statistically significant differences when compared to control (*: p<0.05, paired t-test). Figure 2c: Immunofluorescence analysis of the MERVL-gag protein in control and Foxd3 KO cells. Nuclei are labelled with DAPI. The percentage of counted cells exhibiting the depicted phenotype is indicated. Scale bar represents 10 μm, n=150. Figure 2d: Venn diagram representing the overlap between genes upregulated in Foxd3 KO cells (yellow) and in MERVL: Tomato+ cells (representing 2CLCs, blue) (Macfarlan *et al*, 2012). Figure 2e: Immunofluorescence analysis of 2CLC markers ZSCAN4 and DPPA3 in control and Foxd3 KO cells. Nuclei are labelled with DAPI. The percentage of counted cells exhibiting the depicted phenotype is indicated. Scale bar represents 10 μm, n=150. Figure 2f: Heat map depicting the comparative gene expression profiles of the top 100 upregulated genes in Foxd3 KO cells. The columns indicate expression in Foxd3 KO cells, 2-cell stage embryos and MERVL:Tomato+ cells.

### Immunofluorescence

5 ×10^4^ ES cells were attached to gelatin coated glass slides by using cytospin (Thermo Scientific). Cells were fixed in 4% para-formaldehyde for 10 minutes at room temperature, washed three times with 1X PBS and permeabilized in a 0.05% triton-X solution for 5 minutes. Permeabilized cells were washed twice with 1X PBS and incubated in blocking solution (PBS/0.25% BSA/0.1% Tween-20% and 10% normal goat serum) for 1 hour at room temperature. The slides were incubated with primary antibodies overnight at 4°C. Slides were then washed with 1X PBS and incubated for 1 hour at room temperature with appropriate fluorescently labelled secondary antibody. After washing with 1XPBS, slides were mounted with VECTASHIELD mounting medium containing DAPI. Cells were observed using a Confocal microscope (Zeiss Observer Z1) to detect the fluorescent signal. Images were captured at 63X magnification, analysed with Zen software 2011 SP3 (Black version) and depicted as ‘maximum intensity’ projections from Z stacks of representative ES cells (n=150). Final assembly of images was done using ImageJ the primary antibodies used are as follows: ZSCAN4, Millipore, Catalog number AB4340, 1:250. MERVL-gag, Hangzhou HuaAn Biotechnology, Catalog number R1501-2, 1:250. DPPA3, Abcam, Catalog number ab19878, 1:250.

### Immunoprecipitation

To detect the interaction between FOXD3 and SUV39H1 and H2, GFP-SUV39H1 and GFP-SUV39H2 expressing *Suv39h dn* ES cells were used and to detect interaction between FOXD3 and SETDB1, GFP-FOXD3 expressing Foxd3 cKO ES cells were used. Cells were subjected to immunoprecipitation with GFP-trap dynabeads (Chromotek gtdk-20) following the manufacturers’ protocol. The eluted IP samples were immunoblotted with specific antibodies using established protocols (Bulut-Karslioglu *et al*, 2012). The antibodies used for immunoblot are as follows: Foxd3: Merck Millipore, Catalog number AB5687, 1:500, GFP: Invitrogen ,Catalog number A11122, 1:500, Setdb1: ThermoFisher, Catalog number MA5-15722, 1:500.

## Results and discussion

### FOXD3 binds to specific families of repeat elements

Mouse embryonic stem cells are an important developmental model to study the genetic and epigenetic mechanisms regulating the pluripotent state and cell fate transitions. We chose to identify putative TF binding sites in MSR, MERVL-LTR, MERVL-int and LINE 5’UTR sequences as the expression of these elements is critical in early development. We obtained consensus sequences for these repeats from Repbase (Jurka *et al*, 2005) (Figure S1a) and subjected them to the TF binding site prediction tool PROMO (Messeguer *et al*, 2002). The TFs that were predicted with a p value less than 0.05, a dissimilarity percentage less than 2 and were expressed in mouse ES cells based on published data (Bulut-Karslioglu *et al*, 2014) were considered as significant hits. Binding sites for TFs such as TRM1, YY1, SOX2, FOXD3 were predicted in the repeat sequences (Figure 1a, S1a). Interestingly, the 5’UTR of L1mdA, the promoter region for the transcriptionally competent LINE repeat, was largely devoid of TF binding sites. The forkhead transcription factor FOXD3 emerged as a promising candidate as reports indicate that FOXD3 plays a crucial role in maintaining ES cell pluripotency (Hanna *et al*, 2002; Krishnakumar *et al*, 2016; Respuela *et al*, 2016). FOXD3 is expressed in mESCs and has been shown to bind to and recruit chromatin factors such as HDACs, BRG1, LSD1 etc. to their target sites (Krishnakumar *et al*, 2016). Our analysis predicted the presence of the FOXD3 binding site GAATGTTT (Transfac ID T02290) in certain types of repeat elements, specifically, MSR, MERVL-LTR and MERVL-int (Figure 1b). We examined the binding of FOXD3 to MSR and MERVL DNA *in vitro*. Increasing concentrations (0-2.5μM) of the recombinant GST-FOXD3 fusion protein was incubated with double stranded 5’Cy-5 labelled DNA oligonucleotides representing MSR, MERVL-LTR, MERVL-int, L1mdA in addition to *Sox15* (known target for Foxd3 as positive control) (Plank *et al*, 2014) and *Hprt* (negative control). A concentration dependent mobility shift of the DNA-protein complex was seen for Sox*15*, MSR, MERVL-LTR and MERVL-int oligonucleotides but not for L1Mda and *Hprt* oligonucleotides (Figure 1c). To confirm FOXD3 binding to repeat elements in mESCs, we performed Chromatin immunoprecipitation (ChIP) with an antibody against FOXD3 in mESCs. In agreement with our *in vitro* observations, we detected a significant FOXD3 enrichment over MERVL and MSR sequences, but not over L1Mda (Figure 1d). These results indicate that the transcription factor FOXD3 binds to a subset of repeat elements, especially MERVL and MSRs in mouse embryonic stem cells.

### Foxd3 deletion leads to de-repression of MERVL and a subset of 2CLC genes

*Foxd3* null embryos die after implantation and ES cell lines cannot be maintained (Hanna *et al*, 2002; Liu & Labosky, 2008). To understand the effect of Foxd3 on repeat elements in mESCs, we used a conditional gene inactivation approach. *Foxd3fl/fl;Cre-ER* ES cells (Liu & Labosky, 2008); hence forth referred to as Foxd3 cKO cells (A gift from Patricia Labosky) exhibit a tamoxifen-dependent deletion of *Foxd3*, and an almost complete loss of Foxd3 was seen both at transcript and protein level by day 2 of 4-hydroxytamoxifen (4-OHT) treatment (Figure S1b and c). As the cells underwent extensive apoptosis after day 4 of 4-OHT treatment, all the experiments were carried out at day 2. We performed total RNA-Seq analysis of Foxd3 cKO cells in the absence (control) and presence (Foxd3 KO) of 4-OHT as described in materials and methods. Principal component analysis revealed that control and Foxd3 KO cells exhibited significantly distinguishable expression profiles (Figure S1d). As this study was aimed at understanding the role of Foxd3 in repeat regulation, we investigated the expression levels of annotated repeat elements. 38 repeat elements were significantly upregulated and 1 repeat element was significantly downregulated in Foxd3 KO cells. The most significantly upregulated repeats belonged to the ERV superfamily (Figure 2a) and among the ERVs, ERVL-LTR as well as ERVL-int repeat sequences were most highly upregulated (Figure 2a, S1e). We validated these results by RT-qPCR which showed significant upregulation of MERVL-LTR as well as MSR but not L1MdA in Foxd3 KO cells (Figure 2b). While the MSR consensus sequence contained the FOXD3 binding site (Figure 1a) and FOXD3 was found to bind to MSR in our previous experiments (Figure 1c), MSR was not significantly upregulated in our RNA-Seq data analysis. We therefore limited our further analysis to the more robust upregulation of MERVL upon Foxd3 depletion. We addressed the expression of MERVL at protein level by performing immunofluorescence of control and Foxd3 KO cells using an antibody against the MERVL-gag protein We observed that while less than 1% cells expressed MERVL in control cells, this number increased up to 21% in Foxd3 KO (Figure 2c). These results demonstrate that in mouse ES cells FOXD3 represses the MERVL retroviral element and to a lesser extent, major satellite repeats.

In addition to repeat elements, we also conducted comparative gene expression analysis in control and Foxd3 KO cells. Foxd3 KO ES cells exhibited a significant upregulation of 858 genes and downregulation of 413 genes indicating that in these cells, FOXD3 plays primarily a repressive role, in agreement with previous reports (Respuela *et al*, 2016). Functional classification of the genes upregulated in Foxd3 KO cells using DAVID (Huang *et al*, 2009) showed significant (p< 0.001) enrichment of genes involved in the negative regulation of cell differentiation, cell proliferation, transcription and negative regulation of apoptosis; while cell differentiation, neural crest migration and nervous system development genes were enriched in Foxd3 KO downregulated genes (Figure S2a). This is consistent with the role of Foxd3 in stem cell maintenance and neural crest development (Liu & Labosky, 2008; Lukoseviciute *et al*, 2018). We also compared Foxd3 KO upregulated genes with Foxd3 target genes identified by ChIP-Seq described in (Krishnakumar *et al*, 2016) and found that of the 858 genes upregulated in Foxd3 KO, 132 genes were also targets of Foxd3 (Figure S2b, Supplemental table 1) indicating that their regulation may be a result of direct binding and repression by FOXD3. FOXD3 may control the transcription of other genes indirectly by binding to and sequestering activator proteins, acting as a cofactor in repressive complexes or repressing enhancers that control gene expression (Lam *et al*, 2013; Krishnakumar *et al*, 2016; Respuela *et al*, 2016).

As MERVL was the most upregulated repeat element in Foxd3 KO and MERVL activation is a hallmark of the 2-cell embryonic stage, we compared the genes upregulated in Foxd3 KO with genes upregulated in 2CLCs. Macfarlan et al described Mervl:tomato high cells that represent the population of 2CLCs in an ES cell culture (Macfarlan *et al*, 2012). Comparative analysis showed that 51% of genes upregulated in Mervl:tomato cells were also significantly upregulated in Foxd3 KO cells (Figure 2d). These included 2C specific genes of the Zcan4 family, Prame family, Dppa3, Zfp352, as well as Tcstv1 and 3 (Figure 2e,2f,S2c) (Macfarlan *et al*, 2012; Hendrickson *et al*, 2017). Immunofluorescence analysis of ZSCAN4 and DPPA3 revealed that these proteins were barely detectable in control cells, but expressed at high levels in 41% (ZCAN4) and 28% (DPPA3) of Foxd3 KO cells (Figure 2e) suggesting that the balance between ES and 2CLC is shifted towards 2CLC in a Foxd3 KO cell population. We further compared the top 100 upregulated genes in Foxd3 KO with genes expressed in the embryo at the 2-cell stage (Macfarlan *et al*, 2012). We saw that 56% of the top Foxd3 KO upregulated genes overlapped with either the 2C embryo or MERVL: tomato cell expression data (Figure 2f). This indicates that most highly upregulated genes in Foxd3 KO are 2CLC genes underscoring the contribution of Foxd3 in repressing the conversion of mESCs to 2CLC.

Interestingly, our data did not show an upregulation of Dux, which has been described as a positive regulator of the 2CLC state (Hendrickson *et al*, 2017). Foxd3 seems to function in a Dux-independent pathway to repress MERVL, which is consistent with recent reports indicating that Dux expression is dispensable for zygotic gene activation (Chen & Zhang, 2019; Guo *et al*, 2019; Iaco *et al*, 2020). In addition to MERVL, studies have shown that transient activation of LINE1 RNA is required for the normal developmental progression of a 2C embryo (Jachowicz *et al*, 2017). LINE1 acts to repress Dux in mouse ES cells via a KAP1/nucleolin pathway (Percharde *et al*, 2018). In our studies LINE1 expression remained largely unchanged in Foxd3 KO cells, which may maintain the repression of Dux and hence prevent a complete recapitulation of the 2C transcription profile, as 49% of the genes upregulated in the MERVL-tomato high cells did not appear to be Foxd3-dependent (Figure 2d). Repeat element transcription, therefore, seems to be governed by a collaborative action of multiple modes of regulation and warrants comprehensive studies aimed at understanding the combinatorial regulation of repeats and in turn the balance between mESCs and 2CLCS.

### MERVL de-repression depends on direct binding of FOXD3

To determine whether the upregulation of MERVL is a result of direct occupancy of FOXD3 on the repeat sequences, we generated two DNA binding mutants of Foxd3 based on previous reports identifying the amino acids YSY and FVK in the DNA binding domain of FOXA3 as being essential for the DNA binding function of the protein (Clevidence *et al*, 1993). These amino acids were also conserved in mouse FOXD3 (Figure S3a). We generated recombinant GST-tagged wild type FOXD3 as well as FOXD3 mutant proteins carrying YSY→RAD (M1) or FVK→VAM (M2) mutations (Figure S3b and c). Gel shift assays with Foxd3 M1 and M2 proteins indicated that both mutants failed to bind MERVL and MSR labelled oligonucleotides (Figure 3a). To determine whether a DNA binding deficient FOXD3 could affect repeat element expression and gene transcription, we established Foxd3 cKO mESC lines over-expressing GFP-tagged wild type FOXD3, FOXD3 M1 and FOXD3 M2 (Figure S3d). Foxd3 depletion by tamoxifen led to an expected de-repression of MERVL in non-rescued cells. This de-repression of MERVL was significantly weaker in cells overexpressing wild type FOXD3, but not in FOXD3 mutant (M1) expressing cells, which showed a similar up-regulation as Foxd3 KO (Figure 3b). A similar effect was seen for MSRs, but the LINE1 promoter sequence, which was devoid of the FOXD3 binding site, was not up-regulated in any of the conditions. In the absence of the rescue construct, FOXD3 depletion resulted in MERVL-gag protein expression in 18% of the cells. In contrast, in cells overexpressing wild type FOXD3, MERVL-gag was detected in less than 2% of the cells. This rescue was not apparent in FOXD3 M1 expressing cells (Figure 3c). These results indicate that the DNA binding domain of FOXD3 is required for the repression of MERVL and MSR, and suggest a direct regulation of these repeat elements by FOXD3.

**Figure 3:**
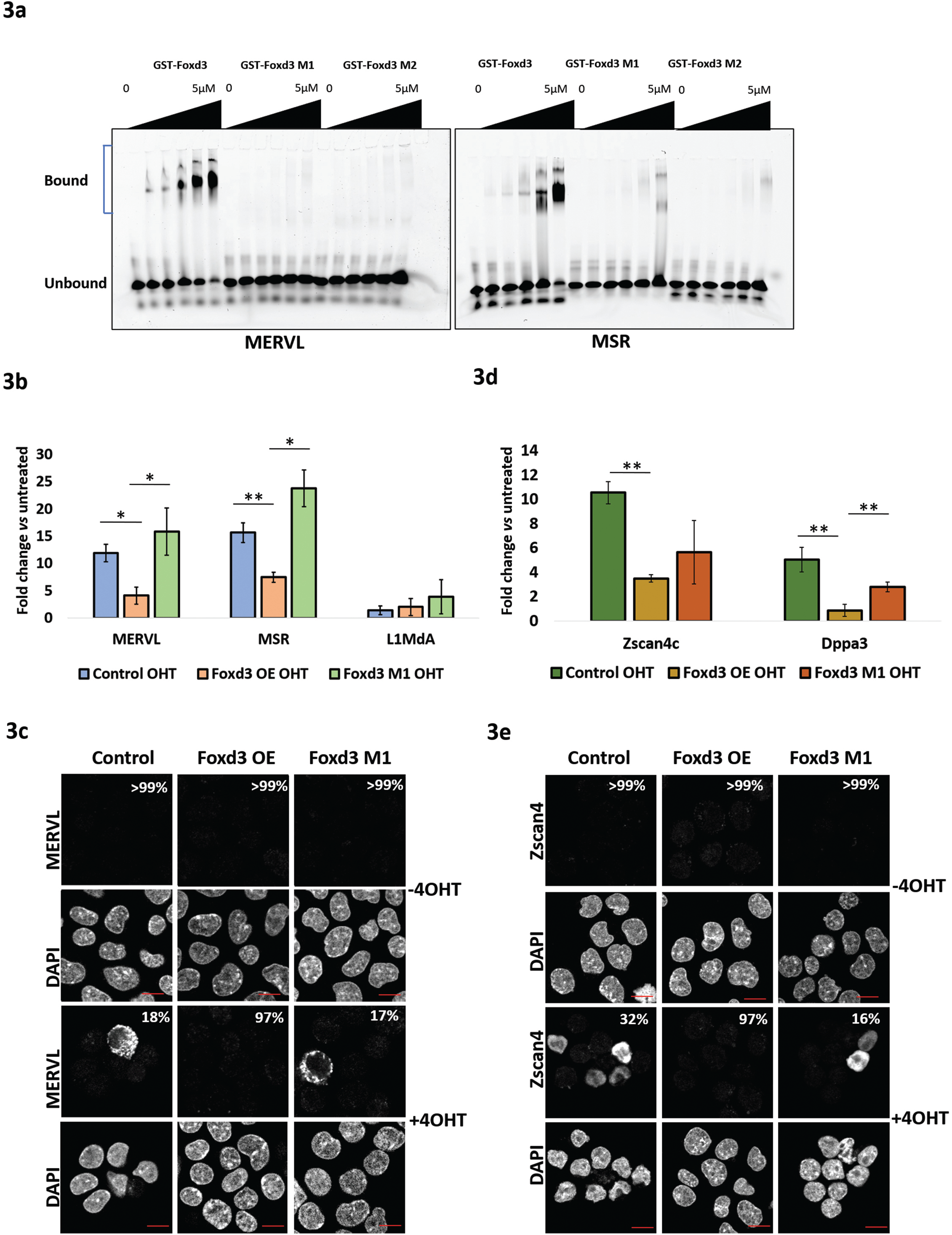
FOXD3 DNA binding domain is essential for regulation of MERVL and 2CLC genes. Figure 3a: Electrophoretic mobility shift assay (EMSA) with increasing concentrations (0, 30nM, 60nM, 1.25μM, 2.5μM and 5μM) of recombinant GST-FOXD3, DNA binding mutants GST-FOXD3 M1 and GST-FOXD3 M2 and fixed concentration (50nM) of 5’-Cy5 labelled dsDNA oligonucleotides (35 bp each) representing MERVL and MSR. Figure 3b: RT-qPCR analysis of MERVL, MSR and L1MdA in Foxd3 cKO cells, Foxd3 cKO cells over expressing full length FOXD3 or FOXD3 M1 without and with 4-OHT. Data is represented as fold change relative to respective untreated cells after normalizing to *Gapdh*. The average data for 2 biological replicates is plotted, error bars represent standard deviation. Asterisks indicate statistically significant differences. (**: p< 0.005, *: p<0.05 paired t-test). Figure 3c: Immunofluorescence analysis of MERVL-gag protein in Foxd3 cKO cells, Foxd3 cKO cells over expressing full length FOXD3 or FOXD3 M1 without and with 4-OHT. Nuclei are labelled with DAPI. The percentage of counted cells exhibiting the depicted phenotype is indicated. Scale bar represents 10μm n=150. Figure 3d: RT-qPCR analysis of, 2C genes *Zscan4c* and *Dppa3* in Foxd3 cKO cells, Foxd3 cKO cells over expressing full length FOXD3 or FOXD3 M1 without and with 4-OHT. Data is represented as fold change relative to respective untreated cells after normalizing to *Gapdh*. The average data for 2 biological replicates is plotted, error bars represent standard deviation. Asterisks indicate statistically significant differences. (**: p< 0.005, *: p<0.05 paired t-test). Figure 3e: Immunofluorescence analysis of the ZSCAN4 protein in Foxd3 cKO cells, Foxd3 cKO cells over expressing full length FOXD3 or FOXD3 M1 without and with 4-OHT. Nuclei are labelled with DAPI. The percentage of counted cells exhibiting the depicted phenotype is indicated. Scale bar represents 10μm n=150.

We investigated whether the differences in MERVL were reflected in the expression of 2C genes. Upon tamoxifen treatment, 2C specific genes such as *Zscan4c* and *Dppa3* showed an upregulation in Foxd3 cKO cells, reduced expression in wild type FOXD3 overexpressing cells and an upregulation in FOXD3 M1 cells (Figure 3d). We also observed a similar result at the protein level where the percentage of FOXD3 depleted cells expressing ZSCAN4 was 32% in the absence of the rescue construct, reduced to 3% in the presence of wild type FOXD3 and increased to 16% in the presence of FOXD3 M1 (Figure 3e), emphasizing the requirement of an intact DNA binding domain of FOXD3 in repressing MERVL and 2C genes

Interestingly, Foxd3 KO led to a higher percentage of ZSCAN4 expressing cells than MERVL expressing cells (Figure 2c, 2e, 3c, 3e). There are conflicting reports regarding the sequence of activation of MERVL and ZSCAN4 proteins during the conversion of mESCs to 2CLCs. Reports indicate that ZSCAN4 activates MERVL, and ZSCAN4 positive cells act as an intermediate state between ES cells and 2CLCs (Zhang *et al*, 2019; Fu *et al*, 2020). In contrast, analysis of transcription dynamics across a pseudo-time trajectory reveals that MERVL expression precedes Zscan4 family gene expression during ESC to 2CLC transition (Eckersley-Maslin *et al*, 2016). Our comparative analysis between Foxd3 binding genes and genes upregulated in Foxd3 KO cells (Figure S2b and Table S1) indicates that Zscan4 family genes are not targets for FOXD3 binding. Activation of Zscan4 in FOXD3 KO cells, therefore, maybe dependent on MERVL expression. Our study leaves open the possibility of a feedback loop between MERVL and Zscan4 family genes that is activated upon MERVL expression in Foxd3 KO cells. It is also possible that the expression of some genes may be independent of the DNA binding activity of FOXD3. Further comprehensive understanding of the transcriptional dynamics may shed more light on the temporal activation and the combinatorial action of gene-repeat expression in the transition from ES to 2CLC state.

### FOXD3 binds to and recruits SUV39H1 to MERVL and MSR

Our data shows that FOXD3 represses MERVL and MSR in mouse ES cells. ERVs and MSRs are largely regulated by the activity of heterochromatin histone methyltransferases such as SETDB1, SETDB2, SUV39H1 and SUV39H2 that establish H3K9me3 to repress transcription (Karimi *et al*, 2011; Bulut-Karslioglu *et al*, 2014; Groh & Schotta, 2017). FOXD3 has been shown to interact with and recruit chromatin modifying proteins to target sites (Respuela *et al*, 2016; Krishnakumar *et al*, 2016). To determine whether FOXD3 could interact with SETDB1/2 or SUV39H1/H2, we performed STRING analysis, which predicted a putative interaction of FOXD3 with SETDB1 and SUV39H1 (Figure 4a). To validate the FOXD3-SUV39H1 interaction in mESCs, we used mESCs devoid of both SUV39H1 and SUV39H2 (*Suv39h dn*) expressing either exogenous GFP-SUV39H1 or GFP-SUV39H2 under the control of a β-actin promoter (Velazquez Camacho *et al*, 2017). We used Foxd3 cKO cells over-expressing GFP-FOXD3 for the FOXD3-SETDB1 interaction. GFP immunoprecipitation followed by immunoblot demonstrated a sub-stochiometric interaction of FOXD3 with SUV39H1 and SUV39H2 (Figure 4b), but none with SETDB1 (Figure S3e).

**Figure 4:**
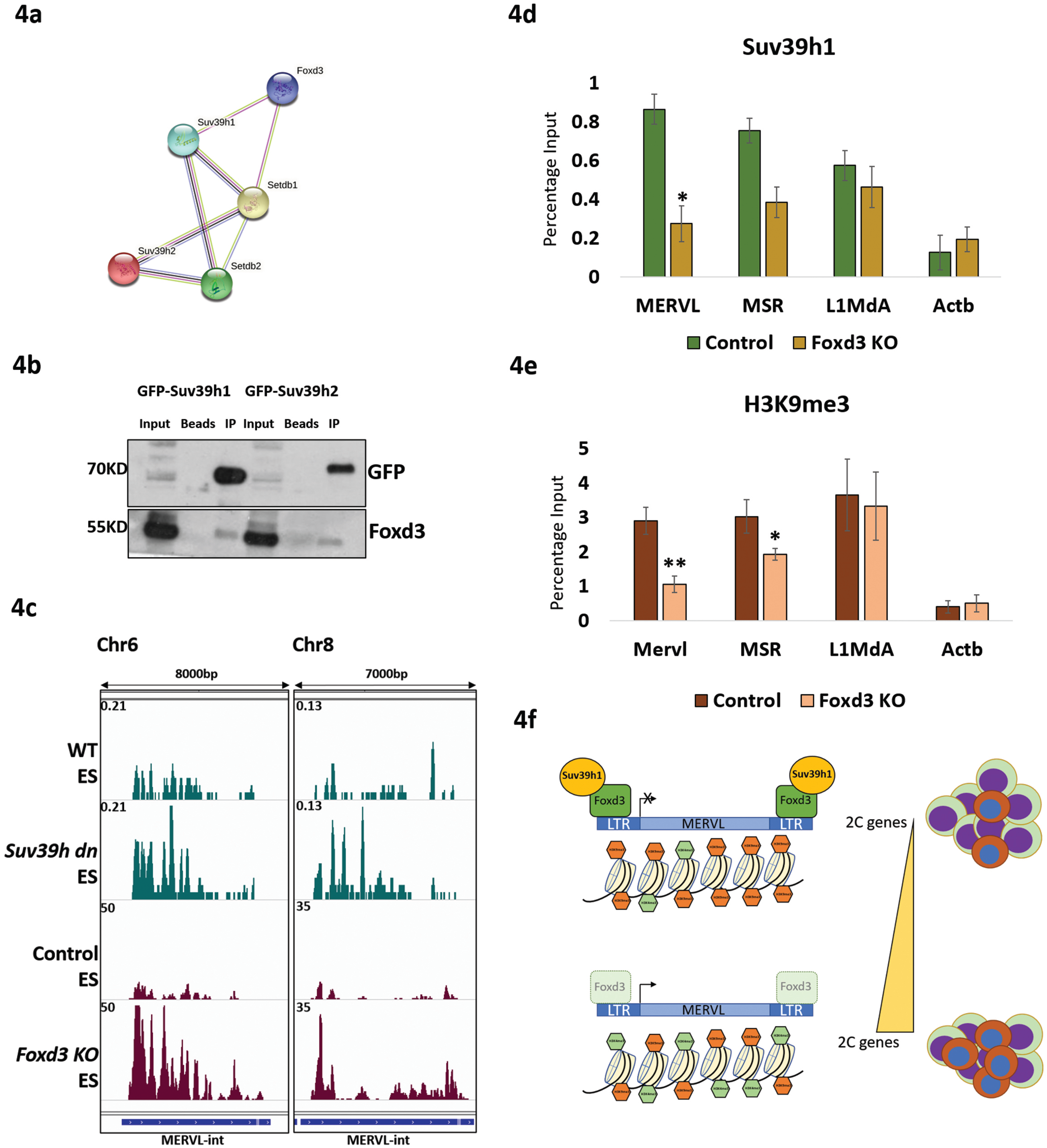
FOXD3 recruits SUV39H1 to repeat elements. Figure 4a: STRING analysis to predict putative interaction between FOXD3, SUV39H1, SUV39H2, SETDB1 and SETDB2. FOXD3 was significantly predicted to interact with SUV39H1 and SETDB1 (p value < 0.01). Figure 4b: GFP immunoprecipitation analysis using GFP-SUV39H1 or GFP-SUV39H2 expressing *Suv39h dn* mESCs. The top panel represents immunoblot using GFP antibody and the bottom panel represents immunoblot using FOXD3 antibody. Figure 4c: MERVL-int expression in WT, *Suv39h dn*, Foxd3 cKO control and Foxd3 KO mESCs as determined by RNA-seq. The browser track shows representative MERVL-int repeat elements. 2 regions encompassing 7000-8000bp from chromosome 6 and chromosome 8 are depicted. Figure 4d: ChIP-qPCR analysis of endogenous SUV39H1 enrichment over MERVL, MSR and L1MdA in control and Foxd3 KO cells. *Actb* promoter is used as negative control. Data is represented as average percentage input from 3 biological replicates. Error bar indicates SEM (*: p<0.05 paired t-test). Figure 4e: ChIP-qPCR analysis revealing H3K9me3 enrichment over MERVL, MSR and L1MdA in control and Foxd3 KO cells. *Actb* promoter is used as negative control. Data is represented as average percentage input of 3 biological replicates. Error bar indicates SEM (**: p<.005, *: p<0.05 paired t-test). Figure 4f: FOXD3 represses MERVL in mouse ES cells: FOXD3 (green) binds to MERVL-LTR and recruits SUV39H1 (yellow) to the LTR. SUV39H1 facilitates repression of MERVL by establishing H3K9me3 (orange hexagons). Foxd3 depletion disrupts the recruitment of SUV39H1 and leads to a reduction in H3K9me3 over MERVL. This in addition to an increase in H3K4me3 (green hexagons) on the region results in the activation of MERVL and a concomitant increase in 2CLC specific genes, shifting the balance between mESCs (Green and purple) and 2CLCs (Brown and blue). Foxd3 thus helps maintain ES cells in a pluripotent state by regulating MERVL.

Bioinformatics analysis of RNA-Seq data from *Suv39h dn* mESCs (Bulut-Karslioglu *et al*, 2014) showed an increase in MERVL-int expression in *Suv39h dn* cells similar to upregulation in Foxd3 KO cells (Figure 4c) indicating that these repeats are at least in part regulated by the SUV39H family proteins. To determine the functional relevance of FOXD3 interaction with SUV39H1/2, we performed ChIP in control and Foxd3 KO cells using an antibody specific for SUV39H1. In control cells, we observed an enrichment of SUV39H1 over MERVL, MSR as well as L1MdA. This enrichment was significantly reduced for MERVL and MSR in Foxd3 KO cells, but not for L1MdA, indicating that FOXD3 participates in recruiting SUV39H1 at these loci (Figure 4d).

Finally, to determine the histone modification profile over MERVL and MSR in Foxd3 KO cells, we performed ChIP for select activating and repressive histone modifications. Our experiments revealed that consistent with the increase in transcription and reduced recruitment of SUV39H1, the H3K9me3 mark decreased over MERVL and MSR repeats in Foxd3 KO cells (Figure 4e). This was also accompanied by a modest increase in H3K4me3 as well as H3K27me3 (Figure S4a). LSD1/Kdm1a, a histone demethylase with catalytic activity towards H3K4, has been shown to repress MERVL in mESCs (Macfarlan *et al*, 2011). Interestingly, LSD1 interacts with FOXD3 in mESCs (Respuela *et al*, 2016). The enrichment of H3K4me3 over MERVL in Foxd3 KO cells may be a consequence of reduced FOXD3-mediated LSD1 occupancy over these repeat regions. We also see an enrichment of H3K27me3 over the repeat regions in Foxd3 KO, which is consistent with previous reports indicating that an increase in H3K27me3 can follow a reduction in H3K9me3 (Peters *et al*, 2003), underscoring the plasticity between these systems. While MSRs, intact ERVs and LINE1 elements are regulated by SUV39H methyltransferases (Bulut-Karslioglu *et al*, 2012, 2014), our data demonstrate that they may be recruited to the repeats using different modes as the SUV39H1 enrichment as well as H3K9me3 levels remain unchanged over the LINE1 promoter in Foxd3 KO cells.

MERVL is known to be repressed by a variety of other regulatory mechanisms. Our data shows that depleting FOXD3 results in a robust upregulation of MERVL in a subset of ES cells and an incomplete recapitulation of the 2CLC transcription profile. Other known activating as well as repressive modulators of MERVL such as Dux or Setdb1 are unchanged in Foxd3 KO. We analysed published data sets (GSE53386) (Fan *et al*, 2015) to determine the expression of Foxd3 through different stages of embryonic development and found that Foxd3 expression begins at the morula stage and is the highest in ES cells (Figure S4b). This suggests that while Foxd3 may not be crucial for the exit from the 2-cell stage, it could facilitate the maintenance of the stem cell state and prevent reversion to the 2CLC state. Combinatorial perturbation of Foxd3 along with other regulatory modules may facilitate a more robust acquisition of the 2CLC transcriptional repertoire. In conclusion, our data demonstrates that in mESCs, FOXD3 binds to and represses MERVL and MSR repeat elements by recruiting the heterochromatin histone methyltransferase SUV39H1 that establishes the H3K9me3 mark to these sites. In the absence of Foxd3, MERVL and MSR repeat elements are de-repressed, and 2CLC genes are activated, skewing the mESC-2CLC balance towards 2CLC (Figure 4f). As repeat element sequences contain numerous other TF binding sites, it is tempting to hypothesise a scenario where a combination of DNA binding proteins act in a context-specific manner to recruit activating or repressive proteins to MERVL and MSRs and regulate their expression in ES cells. It remains to be understood whether a similar transcription factor-mediated regulatory mechanism exists for other repeat elements. This study identifies a novel heterochromatin function for transcription factors and opens up further avenues into the investigation of transcription factor-mediated control of repeat elements in regulating stem cell pluripotency and lineage commitment.

## Acknowledgements

This study was initiated in the lab of Thomas Jenuwein at the Max Planck institute for Immunobiology and Epigenetics (MPI-IE), Freiburg and completed at the National Centre for Cell Science (NCCS) Pune, India. We thank Thomas Jenuwein for lending his support, inputs and expertise in transcription factor-mediated regulation of repeats elements. We also acknowledge Maria Elena Torres-Padilla for insightful discussions regarding MERVL and the 2-cell state. We are grateful to Patricia Labosky for kindly gifting us the Foxd3 cKO cells and Robert Blelloch for arranging the transport of the cells to MPI-IE, Freiburg. We acknowledge the support of the sequencing facility at MPI-IE, Freiburg. We appreciate the help of Gaurishankar Bhaskar in optimising ChIP assays in NCCS, Pune. We thank Deepa Subramanyam for critical reading of the manuscript. Research in D.P. lab is supported by a Department of Science and Technology (DST) INSPIRE faculty award and research in T.J. lab is supported by the Max Planck Society and by additional funds from the German Research Foundation (DFG) within the CRC992 consortium ‘MEDEP’.

## Author contributions

D.P conceptualized, designed and performed the experiments of the project. B.K contributed to the experiments depicted in figures 1c and 3a. B.E and T.M contributed to the experiments depicted in figures 3b and 3c. M.O-S., and D.R performed bioinformatics analysis. D.P wrote the original draft.

## Data availability

The sequencing data generated and reported in this paper can be accessed in GEO using the accession number GSE173602

