## Supplemental information for "Foxd3 controls heterochromatin-mediated silencing of repeat elements in mouse embryonic stem cells and represses the 2-cell transcription program"

#### Supplemental data

##### Figure S1

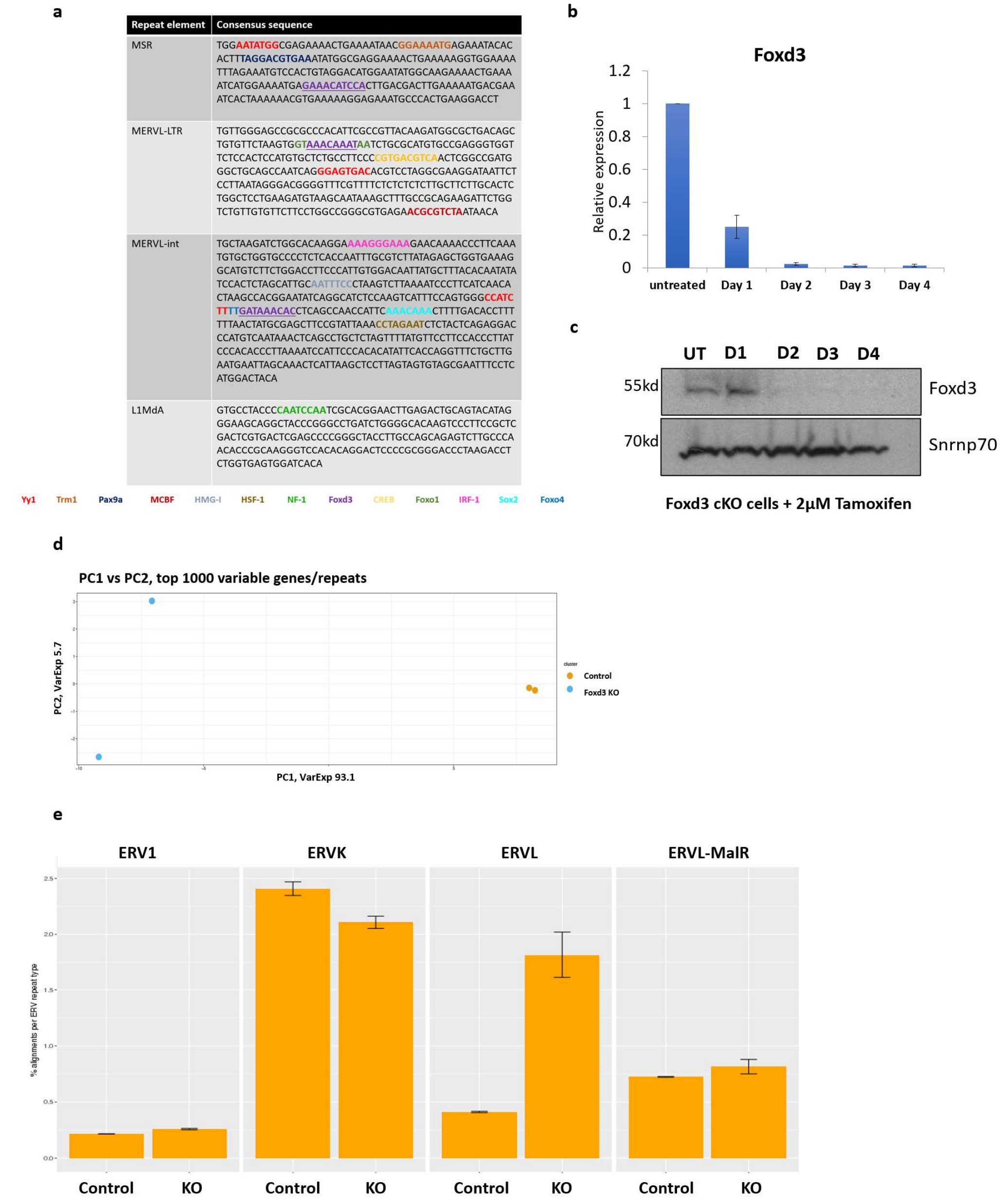

**Supplemental Figure S1:**

S1a: Consensus sequences for MSR, MERVL-LTR, MERVL-int and L1MdA marked with TF binding sites predicted by PROMO. S1b: Foxd3 expression detected by RT-qPCR for untreated and 4-OHT treated Foxd3 cKO cells. Data is plotted as relative expression compared to untreated cells after normalizing to *Gapdh*. S1c: Immunoblot of protein lysates prepared from untreated and 4-OHT treated cells, probed with Foxd3 antibody. Snrnp70 is used as loading control. S1d: Principal Component Analysis of the RNA Seq data from control (Orange) and Foxd3 KO (Blue) indicating percentage variance for 2 principal components. S1e: Expression levels for each ERV subtype plotted along with standard error for control and Foxd3 KO cells.

### Figure S2

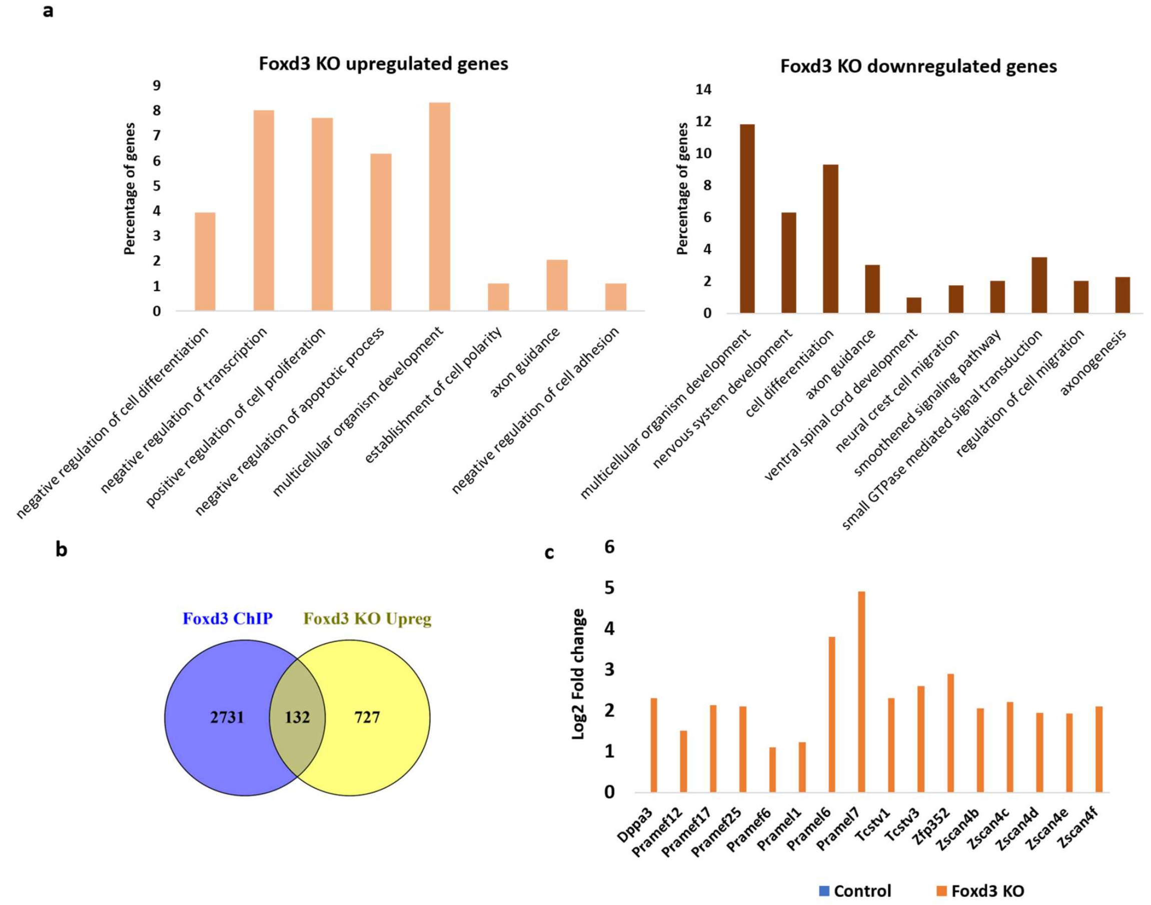

**Supplemental Figure S2:**

S2a: GO protein classes significantly enriched in genes upregulated (left panel) and downregulated (right panel) in Foxd3 KO. The Y axis represents percentage of genes belonging to each protein class. S2b: Venn diagram comparing Foxd3 bound genes (Krishnakumar *et al*, 2016) with genes upregulated in Foxd3 KO cells.S2c: Expression of representative 2CLC genes in Foxd3 KO cells. The Y axis represents log2 fold change in Foxd3 KO cells compared to control cells.

#### Figure S3

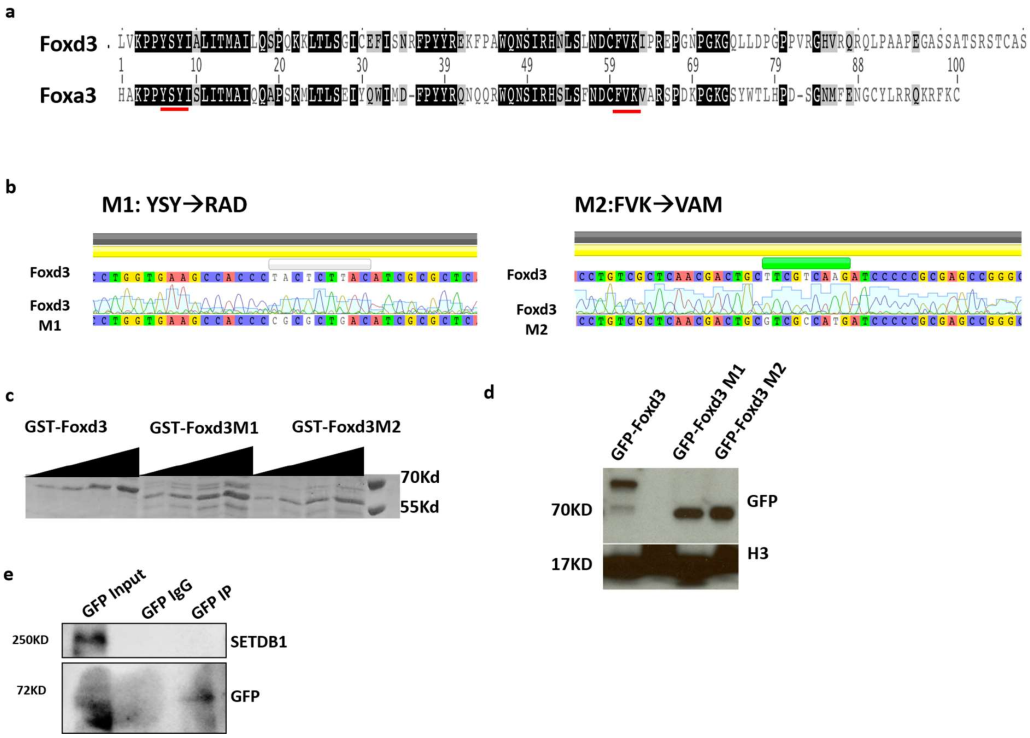

**Supplemental Figure S3:**

S3a: Conservation of YSY and FVK amino acids (underlined) between mouse FOXA3 and FOXD3 DNA binding domain sequences. S3b: DNA sequencing results indicating generation of Foxd3 M1 and M2 mutants. S3c: Coomassie staining indicating expression of recombinant GST-FOXD3, GST-FOXD3 M1 and GST FOXD3 M2. S3d: Immunoblot (α GFP top panel and α H3 bottom panel) for lysates from Foxd3 cKO cells expressing GFP-FOXD3, GFP-FOXD3 M1 and GFP-FOXD3 M2. Histone H3 levels are used as loading control. S3e: GFP immunoprecipitation analysis using Foxd3 cKO mESCs expressing GFP-FOXD3. The top panel represents immunoblot using SETDB1 antibody and the bottom panel represents immunoblot using GFP antibody.

#### Figure S4

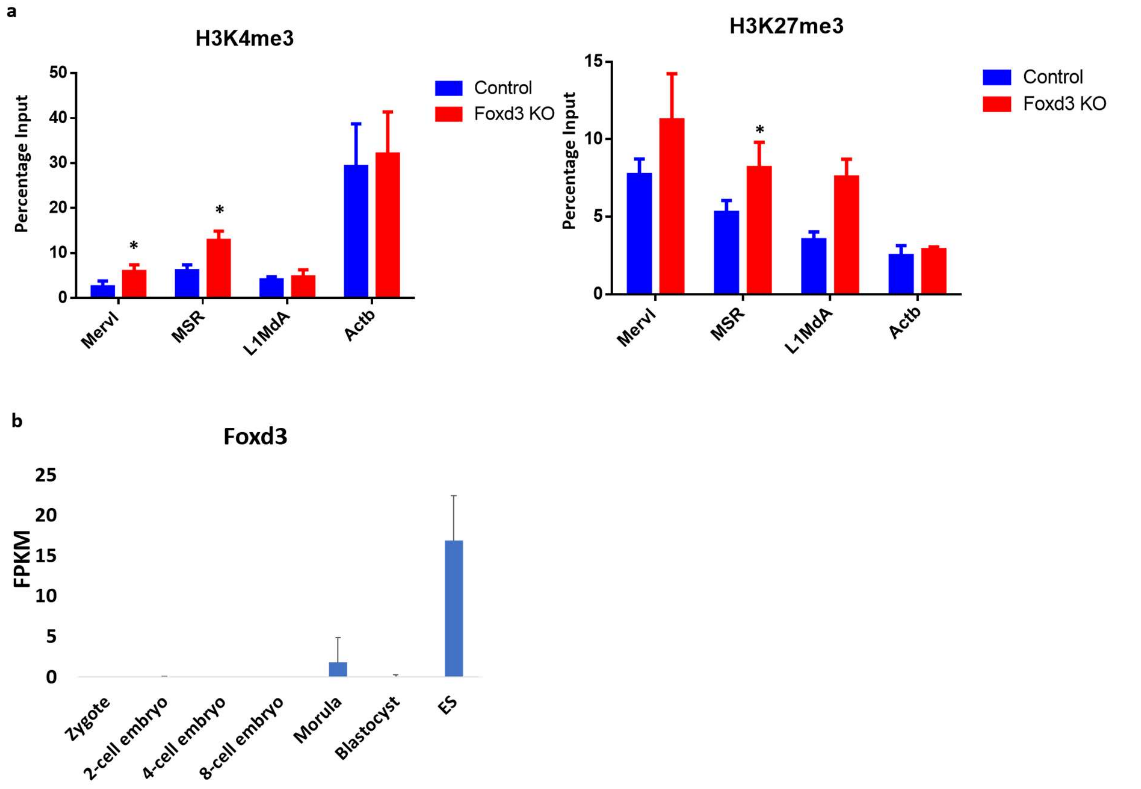

**Supplemental Figure S4**

S4a: ChIP-qPCR analysis for H3K4me3 (left panel) and H3K27me3 (right panel) enrichment over MERVL, MSR and L1MdA in Foxd3 cKO cells without and with 4-OHT treatment. *Actb* promoter is used as negative control. Data is represented as average percentage input of 3 biological replicates. (*: p<.05 paired t-test). S4b: Bar graph depicting Foxd3 expression at different stages of embryonic development. Y axis depicts average FPKM from 4 single cell RNA Seq data sets. Error bar represents standard deviation

**Supplemental table 1: Foxd3 targets upregulated in Foxd3 KO cells**

| | | | |
|---|---|---|---|
| 0610040J01Rik | Fam81a | Lnx1 | Rapgefl1 |
| 2010107G12Rik | Fbxo15 | Lonrf1 | Rbms3 |
| 4930506M07Rik | Fgfbp1 | Lpar1 | Rcor2 |
| 6430573F11Rik | Flnb | Lrpap1 | Rcsd1 |
| Abca4 | Frmpd1 | Lxn | Rimkla |
| Abcb5 | Fundc1 | Mfge8 | Robo3 |
| Adamts7 | Gabra1 | Mobp | Ror1 |
| Adamts8 | Gad2 | Mpped2 | Rragd |
| Aebp2 | Galnt3 | Mreg | S1pr1 |
| Alpl | Gata6 | Nabp1 | Satb2 |
| Alppl2 | Gjb3 | Nanos3 | Sh2d4b |
| Ankrd45 | Glis3 | Nat1 | Slc25a4 |
| Aox3 | Gpd1 | Nelfa | Slc7a9 |
| Ap3b2 | Gprc5c | Ngfr | Slco2a1 |
| Aqp3 | Gprin3 | Nptx2 | Sorbs1 |
| B020004J07Rik | Gsta3 | Nrp2 | Sox21 |
| Bcl2l14 | Hmga2 | Nt5e | Spesp1 |
| Cadps2 | Hmgn5 | Nudt4 | Spic |
| Cbln1 | Id2 | Oasl2 | Tbc1d8 |
| Ccdc17 | Id3 | Otud1 | Tbx3 |
| Ccdc60 | Id4 | Otx2 | Tfap2c |
| Clcn5 | Igf2bp2 | Pcdh19 | Ticrr |
| Col5a2 | Il6ra | Peg12 | Tmcc3 |
| Cpsf4l | Inpp4b | Phf13 | Tmem132c |
| Csf3r | Ipmk | Pkd1l1 | Trpc6 |
| Ctnnal1 | Kcnn2 | Plekhg1 | Ulk1 |
| Dlgap2 | Kifc3 | Podxl | Urgcp |
| Dppa2 | Klf2 | Popdc3 | Veph1 |
| Dppa3 | Lax1 | Pou6f2 | Wtap |
| Dysf | Lbh | Pramel6 | Yes1 |
| Egfl6 | Lmo7 | Prep | Zbtb10 |
| Eomes | Lmx1a | Prickle1 | Zdhhc23 |
| Eps8l2 | Lmx1b | Rab11fip4 | Zfp560 |

**Supplemental table 2: Primers used in the study**

| Name | Sequence | Experiment |
|---|---|---|
| Mervl-LTR | AGACTCAAAGCAAAACAGGCTCCTAGAGGGGAGGT | EMSA |
| MERVL-int | CCATCTTTTGATAAACACCTCAGCCAACCATTCAA | EMSA |
| MSR | CTGAAAATCATGGAAAATGAGAAACATCCACTTGA | EMSA |
| Sox15 | CAGAGGCTACTCTGAAACAAATAAAGAGATATAAA | EMSA |
| L1MdA | GTACATAGGGAAGCAGGCTACCCGGGCCTGATCTG | EMSA |
| HPRT | AGGGCGGGCCGAGGGGCGGAGCCTGGCCGGCAGCG | EMSA |
| Major Sat - F | TGGAATATGGCGAGAAAACTG | RT-qPCR, ChIP |
| Major Sat - R | AGGTCCTTCAGTGGGCATTT | RT-qPCR, ChIP |
| MERVL-F | TTTCTCAAGGCCCACCAATAGT | RT-qPCR, ChIP |
| MERVL-R | GACACCTTTTTAACTATGCGAGC | RT-qPCR, ChIP |
| L1_promoter - F | ACTGCGGTACATAGGGAAGC | RT-qPCR, ChIP |
| L1_promoter - R | TGTGATCCACTCACCAGAGG | RT-qPCR, ChIP |
| Sox15_promoter-F | GGTGCTTGAGAATTGAGACA | ChIP |
| Sox15_promoter-R | GTCTTGTTCTTCCCAGTCCT | ChIP |
| Actb-F | AGCCAACTTTACGCCTAGCGT | ChIP |
| Actb-R | TCTCAAGATGGACCTAATACGGC | ChIP |
| Foxd3-F | GTCCGCTGGGAATAACTTTCCGTA | RT-qPCR |
| Foxd3-R | ATGTACAAAGAATGTCCCTCCCACCC | RT-qPCR |
| IAP-F | GCACCCTCAAAGCCTATCTTA | RT-qPCR |
| IAP-R | TCCCTTGGTCAGTCTGGATTT | RT-qPCR |
| Zscan4-F | CCTCCCTGGGCTTCTTGGCAT | RT-qPCR |
| Zscan4-R | AGCTGCCAACCAGAAAGACACTGT | RT-qPCR |
| Dppa3-F | CGGGGTTTAGGGTTAGCTTT | RT-qPCR |
| Dppa3-R | GGACCCTGAAACTCCTCAGA | RT-qPCR |
